# Effects of temperature gradient on flower and fruit traits: a meta analysis

**DOI:** 10.64898/2026.08.06.743200

**Authors:** Omer Nevo, Evangelia Linda Chronopoulou, Anna E. Azeroth, Demetra Rakosy, Renske E. Onstein, Tiffany M. Knight, Jonas Kuppler

**Author notes:** Equal contribution (Omer Nevo, Jonas Kuppler).

## Abstract

Pollination and seed dispersal by animals are key drivers of terrestrial biodiversity and ecosystem functioning. Effective mutualistic interactions rely heavily on temporal and functional-trait matching between plants and animals. While global warming is known to induce shifts in plant traits, the extent and direction in which higher temperatures may systematically alter flower and fruit traits across species in natural habitats remains poorly understood. Using elevation as a proxy for temperature across natural populations, we conducted a meta-analysis evaluating 21 quantitative functional traits (15 floral, 6 fruit) across 82 studies and 161 species. Standardized mixed-effects linear regression models revealed widespread, systemic responses to elevational temperature gradients in both reproductive structures. In flowers, higher temperatures were systematically associated with changes in petal and sepal width and length (and hence morphology), longevity, nectar volume, number of flowers, and inflorescence length. In fruits, elevation was associated with changes in vitamin C content, crop size, weight and width. Taken together, these results demonstrate that warming temperatures exert widespread, multi-axis effects on the morphology, availability, timing, and nutritional quality of both flowers and fleshy fruits. Given that flower and fruit traits are developmentally linked and co-determine animal visitor dynamics, these temperature-driven phenotypic shifts are likely to propagate cascading disruptions throughout plant–pollinator and plant–frugivore interaction networks under continued climate change.

## INTRODUCTION

Angiosperms - flowering plants - first appeared during the Cretaceous and rapidly reached ecological dominance (Friis et al., 2010). This is often attributed to the two key inventions in their common ancestor, flowers and fruits, and to their their ability to repeatedly and flexibly change functional traits that allow them to survive, adapt, and diversify at higher rates than other plants (Crepet and Niklas, 2009; Onstein, 2020). Flowers and fruits facilitate plant reproduction by supporting the gamete production, promoting pollen exchange, protecting the developing seed and enabling dispersal and thus driving adaptation, diversification, and adaptive radiation (Specht and Bartlett, 2009; Sauquet et al., 2017; Pereira and Coimbra, 2019). While not universal, a central strategy in both pollination and seed-dispersal stages is the attraction of animal mutualists. Pollination by animals allows efficient and directed transfer of pollen among conspecifics, even at low population densities and when distances among individuals are relatively long (Regal, 1982); and biotic seed dispersal allows plants to spread seeds over long distances (Beckman and Sullivan, 2023) while maintaining relatively large and hence energy-dense seeds that offer a competitive advantage in light-poor environments (Bolmgren and Eriksson, 2005). As such, animals play a major role in both stages of angiosperm reproduction, where up to 90% of species rely on animal pollinators (Ollerton et al., 2011; Tong et al., 2023), and around 40% of temperate and up to 90% of tropical woody species depend on seed dispersal by animals (Jordano, 2000). As a result, flowers and fruits are also a major resource supporting hundreds of thousands of animal species and countless more communities (Daniel Kissling et al., 2009; Pineda-Munoz and Alroy, 2014; Ollerton, 2017). The nectar, pollen, and nutrients in flowers and fleshy fruits thus form a major part of a key ecological niche and a major factor determining the structure of the adaptive landscape in which animals live and evolve.

As such, pollination and seed dispersal interaction networks are a central component of many, if not the majority, of terrestrial ecological systems. The structures of these networks are complex, showing diverging degrees of specialization, asymmetry, connectance, heterogeneity, modularity, and nestedness (Bascompte and Jordano, 2007; Olesen et al., 2007; Vázquez et al., 2009; Guimarães et al., 2011). This complexity results from constrained interactions, where each species interacts with only a subset of the potential partners available in the community.

Realized pollination and seed-dispersal interactions require both a temporal and functional-trait matching between the plant and the animal (Garibaldi et al., 2015; McFadden et al., 2022; Guerra et al., 2025). For example, flowers that bloom at night will interact with nocturnal but not diurnal pollinators (Bloch et al., 2017); or flowers that bloom and fruit which mature during a certain of the year will only be able to interact with animals active around this time (Forrest, 2015; Gérard et al., 2020). In addition, both animals and plants show tremendous functional trait diversity across multiple axes: morphological, chemical, physiological, and sensory. Realized interactions depend on trait matching, which ensures that the interaction is possible and that it provides nutrients to the animal and pollination or seed dispersal to the plant. For plant-pollinator interactions, matching between the size of floral structures and the length of pollinator feeding structures can determine which interactions are possible; for example, exceptionally long corolla tubes may be accessible only to pollinators with correspondingly long tongues (Muchhala, 2006). Similarly, fruit and seed size are major determinants of interaction with frugivores who either avoid fruits that are too big (Galetti et al., 2013; Guerra et al., 2025) or rather prefer larger fruits (Valenta et al., 2020). Floral and fruit pigmentation, which gives many of them their familiar conspicuous displays, is functional only when the animal mutualist on the other side has the capacity to perceive color (Nevo et al., 2018b; Valenta et al., 2018; van der Kooi et al., 2018), and scent signals are primarily present and effective in flowers and fruits that interact with animals which tend to rely on their sense of smell (Dobson, 2006; Nevo et al., 2018a; Santana et al., 2021). Thus, pollination and seed-dispersal networks and the ecological systems around them depend heavily on a temporal and functional-trait matching between animals and plants.

This trait matching is usually studied at the species level: species are assumed to be more or less uniform, either allowing an interaction to take place or not. But individuals and populations also differ markedly *within species*. Within species, flowers can vary in scent chemistry (Kuppler et al., 2016; Delle-Vedove et al., 2017), morphology (Kuppler et al., 2016), color (Kellenberger et al., 2019), and phenology (Kuppler et al., 2016). Similarly, fruits of the same species vary in size (Galetti et al., 2013), chemical profiles (Gelambi and Whitehead, 2023; Nguyen et al., 2025; Meyer and Nevo, 2026), and color (Traveset and Willson, 1998; Gómez-Devia and Nevo, 2024). While some of this variation is genetic (Ogundiwin et al., 2009; Kellenberger et al., 2019), a substantial proportion of intraspecific variation in flower and fruit traits is plastic and derives from responses to environmental factors such as temperature, sunlight conditions, water and nutrient availability, and biotic conditions (Marsh et al., 1996; Camargo et al., 2017; Campbell et al., 2019; Karl and Peck, 2022). This means that the probability of trait matching and hence successful pollen transfer or seed dispersal is not constant among each plant-animal species pair: even among species that in theory are expected to interact, some individuals will bear traits closer to the optimum required by their mutualist and are therefore more likely to be attractive to them; while others, closer to the extreme, will be less favorable or even inaccessible.

The substantial environmentally-driven intraspecific variation in flower and fruit traits highlights the vulnerability of these interaction networks to ongoing global warming (Nowak et al., 2022). Earth has already warmed by approximately 1° C above pre-industrial levels and is projected to experience significantly more warming even under the most optimistic scenarios (Intergovernmental Panel on Climate Change (IPCC), 2023). Global warming brings about potential direct effects: higher temperatures may affect functional flower and fruit traits. But it is also likely to have multiple indirect effects, through changes in precipitation patterns to changes in the biotic environment. Indeed, multiple studies from the past years indicate the potential susceptibility of flower and fruit traits to global warming. Physiological changes in plants in response to heat and reduced precipitation can result in changing floral traits (Scaven and Rafferty, 2013; Borghi et al., 2019; Descamps et al., 2021), such as decreases in flower size, reduced nectar production (Mu et al., 2015), phenological changes in flowering time (Gérard et al., 2020; Nicholson and Egan, 2020; Kuppler and Kotowska, 2021; Martén-Rodríguez et al., 2025), and reduced longevity (Blionis and Vokou, 2001; Fabbro and Korner, 2004; Pacheco et al., 2016; Arroyo et al., 2017). The strength or effect size of these effects may vary between species, study and environment. This may result in variable susceptibility of different traits to environmental changes such as global warming (Kuppler and Kotowska, 2021; Day Briggs and Anderson, 2025; Martén-Rodríguez et al., 2025). Other studies, using elevational gradients as proxies for temperature, did not find systemic effects of climate change on flower traits (Novaes et al., 2024). Importantly, these effects have been largely studied in experimental settings and the corresponding patterns in natural settings could further verify the occurrence of the anticipated changes. Moreover, many of those results are from studies that record temperature (or elevation) as a binary character (present/absent; lower/higher), which makes it difficult to extrapolate into the future (i.e. the effect of additional warming). Data for fruits are significantly more scarce and derive primarily from agriculture, but also indicates that warming might change various fruit traits by lowering size, hardness, and nutrient content, and increasing volatile emission rate (Welles and Buitelaar, 1988; Nerd et al., 1999; Fischer et al., 2022; Gómez-Devia and Nevo, 2024).

As such, there is growing evidence that flower, and potentially also fruit, traits may be affected by climate change. However, the degree to which warming systematically alters either fruits or flowers remains unclear. Moreover, flowers and fruits are interconnected in multiple ways: not only do they show ecological similarities - they are also developmentally linked since fruits develop directly from flowers. Thus, in addition to direct effects (e.g. changes in water content due to increased heat), fruit traits can also be altered as a result of changes to floral traits. For example, lower pollination quality can lead to decreased fruit quality and hence reduced seed dispersal capacity (Wietzke et al., 2018). Yet despite the ecological similarities and developmental links, the potential effects of climate change on flower and fruit traits are rarely addressed together.

The goal of the current study is to assess to what degree global warming may drive systemic and across-species change in a broad array of flower and fruit functional traits to help assess whether climate change may alter pollination and seed-dispersal networks by changing floral and fruit traits. We use the elevation-for-temperature approach, in which genetically-continuous populations that grow across an elevational gradient naturally experience different temperature regimes. We conducted a meta analysis in which we systematically extracted and analyzed trait data from 82 studies (71 for flowers; 11 fruits) and 161 species (149 flowers; 12 fruits). We cover all traits in which warming may drive changes in either temporal or trait matching, focusing on those for which sufficient data was available (Figures 1, 2). We use mixed-effects models to assess whether any of the traits show a systemic trend across elevational gradients.

**Figure 1.**
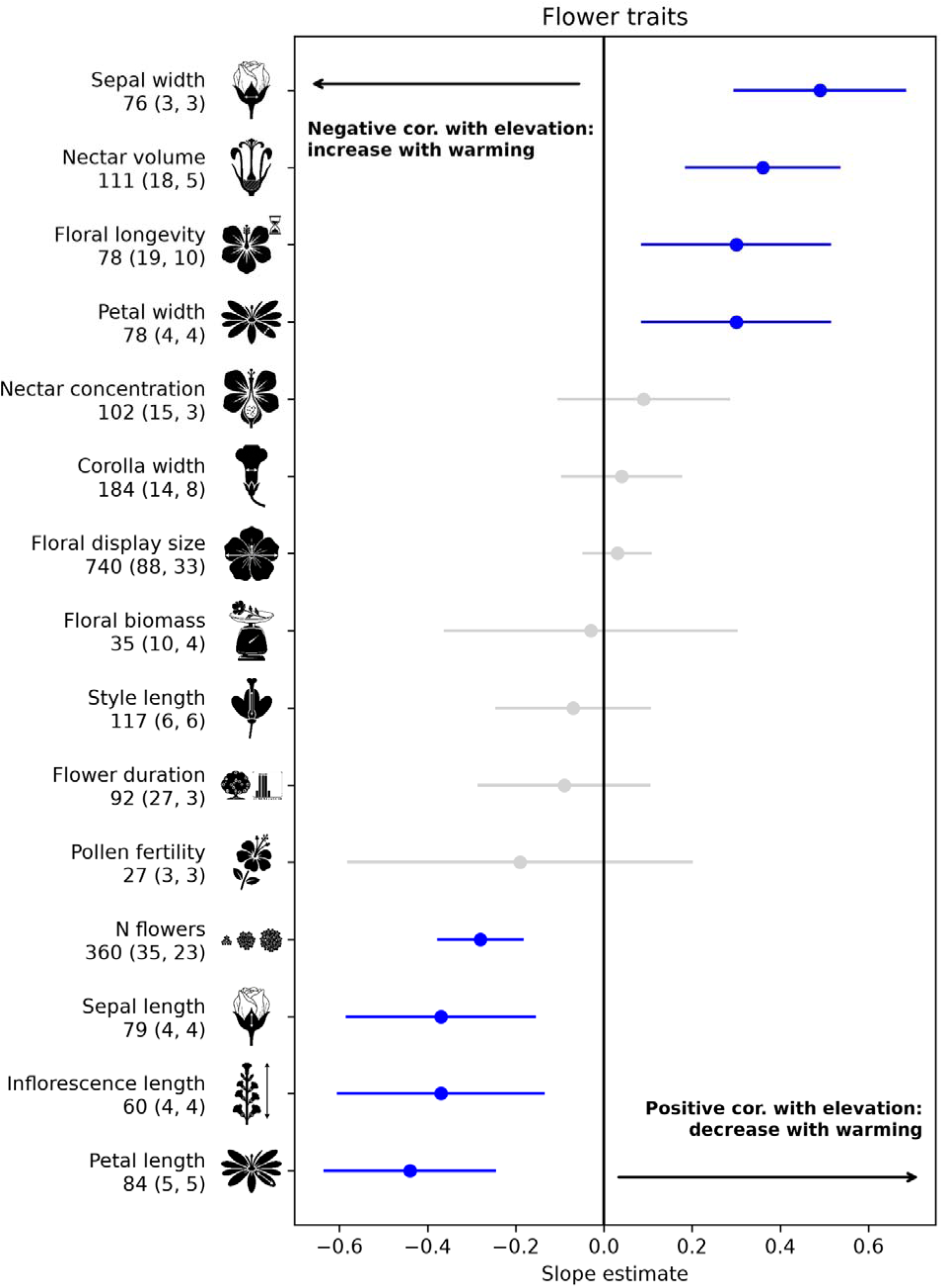
The relationship of floral traits and elevation - model outputs. Each line is a single model for a single trait. Dot represents the model slope estimate. Bars indicate 95% confidence interval. Blue: statistically significant at alpha < 0.05. Traits with a positive slope (right of the black vertical line at 0) are positively correlated with elevation, i.e. negatively correlated with temperature, while those to its left are negatively correlated with elevation and hence positively with temperature. Numbers indicate the number of datapoints (number of species, number of studies).

## METHODS

### Dataset

The Web of Science (ISI) database (Indexes = Web of Science Core Collection, BIOSIS Citation Index, Current Content Connect, Zoological Record, IC Timespan = All years) was searched in July 2021. The search string was composed of two substring combinations: Each combination contained one substring targeting the intervention (elevational or temperature gradient), and one targeting the outcome:

Search term = (“temperatur* gradient” OR “elevation*” OR “altitud*” OR “thermal* gradient” OR “climate gradient” OR “climatic gradient” OR “mountain” OR “alpine”) AND either

Search term = (“floral trait*” OR “flower trait*” OR “floral characteristic*” OR “flower characteristic*” OR “floral volatile*” OR “flower volatile*” OR “floral phenology” OR “flower phenology” OR “flower reward*” OR “floral reward*” OR “floral resource*” OR “flower resource*” OR “food reward*” OR “nectar production” OR “nectar amount*” OR “nectar volume*” OR “nectar” OR “pollen production” OR “pollen amount*” OR “pollen grain*” OR “floral morpholog*” OR “flower morpholog*” OR “floral display*”OR “flower display*” OR “flower size*” OR “floral size*” OR “flower display size*” OR “floral display size*” OR “floral diameter*” OR “flower diameter*” OR “inflorescence size*” OR “inflorescence display size*” OR “inflorescence diameter*” OR “floral shape*” OR “flower shape*” OR “flower symmetr*” OR “floral symmetr*” OR “tube length” OR “tube width” OR “tube opening” OR “corolla*” OR “nectar tube*” OR “petal*” OR “pistil*” OR “stamen*” OR “anther*” OR “calyx*” OR “onset of flower” OR “number of flower*” OR “flowering length”)

Search term = (“fruit characteristic*” OR “fruit trait*” OR “fruit morphology” OR “fruit size” OR “fruit mass” OR “fruit weight” OR “fruit length” OR “fruit diameter” OR “fruit volume” OR “fruit nutrient content” OR “sugar content” OR “fat content” OR “protein content” OR “fiber content” OR “nitrogen content” OR “fruit* phenology” OR “fruit* period” OR “fruit* length” OR “fruit number” OR “number of fruit*” OR “crop size” OR “crop mass” OR “crop weight” OR “crop number”).

### Study selection

The initial search for Web of Science found 3005 publications for flowers (Table S1) and 4046 publications for fruits (Table S4). At a later stage we added 12 additional publications on flowers, which were identified to fit the same criteria despite not having been automatically detected. First, all publications were screened based on title and abstract using the *abstract_screener*()-function of the *metagear*-package (Lajeunesse, 2016). At this stage the evaluation was based on the information provided in the title and the abstract when the abstract was automatically accessed. The evaluation decision fell under 3 categories answering the question *“Should this publication be included based on the abstract?”, “YES”, “MAYBE”, and “NO”.* There were cases that the abstract could not be automatically accessed and as such the evaluation was done based on the title alone. In case of uncertainty the decision was made as “MAYBE” minimizing cases of false negatives. For both searches, we selected all studies that clearly met, or seem to meet, the following criteria (1) any angiosperm plant species (2) any type of elevational and/or temperature gradient, (3) animal pollinated/dispersed species, and (4) measurable of floral or fruit traits relevant for the interaction with animals in the same plant species across at least two locations. After this screening, all publications marked YES or MAYBE passed to the next phase in which the full text was read: 508 publications for floral traits (Table S2) and 187 for fruits traits (Table S5). At this stage, if we could not access the full text of a publication in any other way, we contacted the corresponding author once to acquire the full publication or the raw data of interest. If no response was received by the end of November 2024 the paper was excluded from the study. The top exclusion reasons in both cases included lack of suitable traits, lack of elevational gradients, human activity affecting the studied population and failure to access the necessary dataset. All exclusion criteria are available in Sup tables 1-2,4-5.

### Data collection and extraction

For each study, we extracted the mean, variation (standard error or deviation) and sample size for each population/location separately. Publications fitting all criteria but lacking variation measures were excluded unless the authors provided the full data as they were not considered statistically robust. If a study included multiple distinct elevational levels, all data were extracted. In cases a broad elevational range was provided but not specific information regarding the exact elevation the data points were collected from, the publication was excluded. Data were extracted from tables, texts or graphics. When relevant data were not available online or at institutional libraries, we contacted the corresponding author once and requested the additional data. Data from graphics were extracted using WebPlotDigitizer to the best possible accuracy (https://automeris.io/WebPlotDigitizer/). For measures of variation, the lower and upper error bar, if available, were extracted and the mean value was taken. Extracted results and sources are available from the published data files.Floral and fruit trait names that described the same functional trait but were not consistently named (e.g. N flowers, number of flowers) were consolidated at this stage; some later consolidation followed (see “statistical analysis”).

Elevational gradients were extracted for each measurement location as well as the temperature and GPS-coordinates if available. During the evaluation of the publications 12 additional papers were manually identified and included in the study and data was extracted, same as before the author was contacted once to provide the necessary data and if no response was received the paper was excluded. In total, we were able to extract data out of 119 studies including 237 plant species from 88 genera and 52 plant families for floral traits (Table S3) and 11 studies including 12 species from 10 genera and 9 plant families for fruit traits (Table S6). The results thus reflect the status of the knowledge at the search timepoint, while the inclusion of further species under various conditions potentially may result in different overall patterns and different patterns between species with varying ecology. The final datasets can be found in Tables S3,S6; all further processing was conducted in the script described below.

### Statistical analysis

All analysis of the final extracted dataset was conducted on Python (Python Core Team, 2023) using libraries Pandas (McKinney, 2010), NumPy (Harris et al., 2020), matplotlib (Hunter, 2007), statsmodels (Seabold and Perktold, 2010), scipy (Virtanen et al., 2020), os, contextlib, openpyxl. The script first corrected some variable names that were inconsistent with the dataset or recorded at the wrong column. It then consolidated response variables that were recorded under a different category but were functionally identical (e.g. “floral display size” and “corolla diameter”). The mapping was conducted for fruits and flowers separately using excel tables that were loaded into the code and are available at the GitHub repository for reproducibility. We then removed all datapoints for species that were measured only once (i.e. fewer than the minimum two datapoints per species within a study across an elevational gradient).

We then filtered the datasets to include only datapoints where, per response variable (= trait; e.g. “petal length”), there were at least 25 datapoints from at least three species across at least three studies. We included only quantitative traits. This yielded 15 traits from 149 species (38 families) in flowers, and 6 traits from 12 species (9 families) in fruits (Figures 1,2) across broad elevational gradients from 0 to 5500 m above sea level (Figures S1-S2).

**Figure 2.**
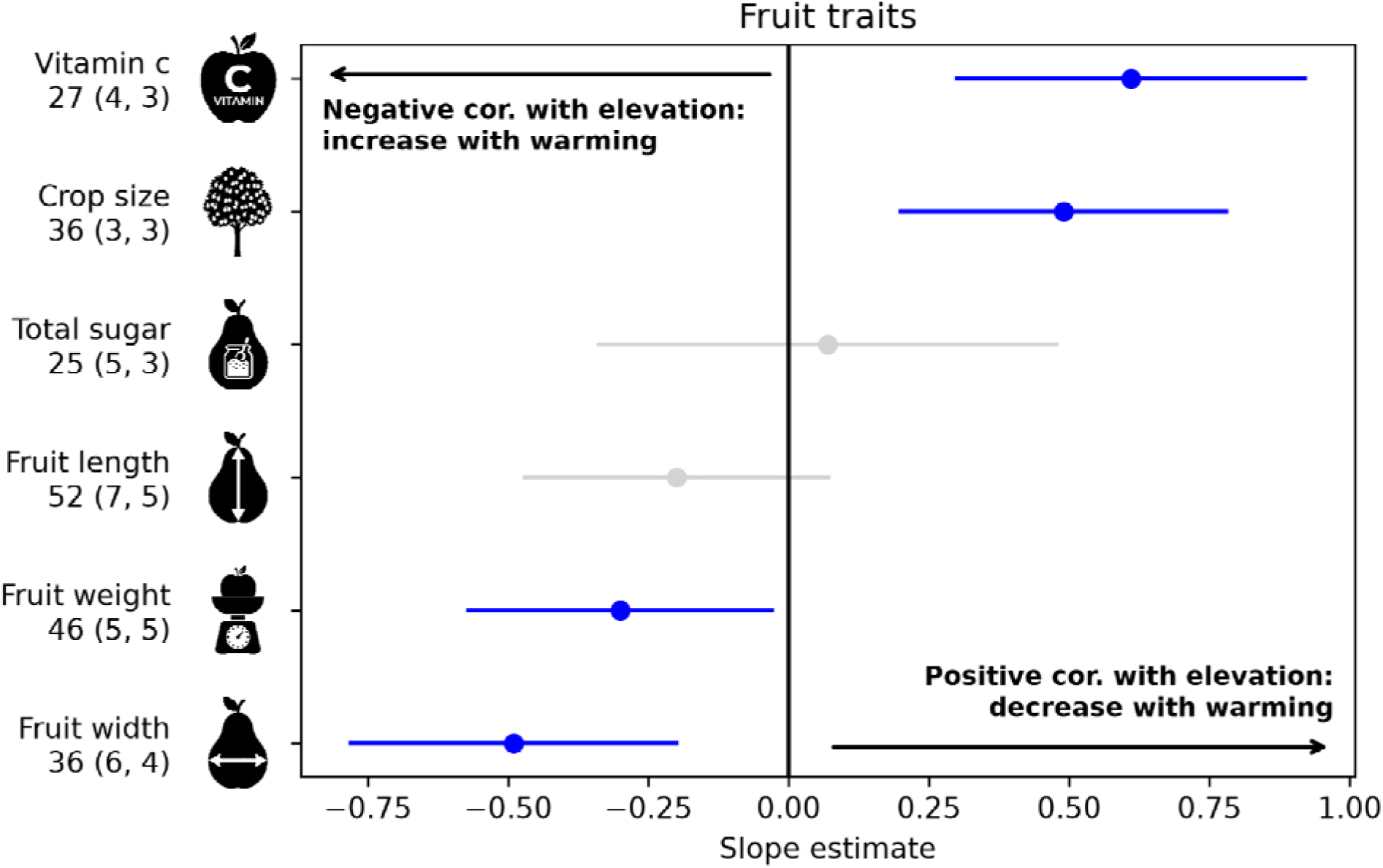
The relationship of fruit traits and elevation - model outputs. Each line is a single model for a single trait. Dot represents the model slope estimate. Bars indicate 95% confidence interval. Blue: statistically significant at alpha < 0.05. Traits with a positive slope (right of the black vertical line at 0) are positively correlated with elevation, i.e. negatively correlated with temperature, while those to its left are negatively correlated with elevation and hence positively with temperature. Numbers indicate the number of datapoints (number of species, number of studies).

The objective of the study was to identify general patterns in the response of flower and fruit traits to variation in temperature within their natural habitat. To allow this comparability, we normalized (=z-transformed) all data (elevation and trait value) *within a species within a study*. As such, the following analyses do not measure, for example, whether an increase in 50 m changes fruit length by a certain number of centimeters, as this will not be comparable across species. Instead, the standardization allows the statistical models to test whether, for example, an increase in one SD in the elevational gradient *within the natural range of the population* is associated with an increase of X SDs in the size of the fruits *of that species*. We visually examined the resulting z-transformed variables to verify that no further transformation was required (available in the jupyter notebook in the GitHub repository).

Data were analyzed per trait separately. In all, the general approach was a mixed effects linear regression model where the z-transformed trait value variable was the response variable, and z-transformed elevation was the sole predictor. In all analyses we included a random intercept factor to account for random variation among the studies and species. The exact structure of the random effect was determined by the structure of the data. We used (a) a simple single factor random effect when each study included exactly one species, and species did not repeat across studies; (b) a nested structure (species within study) when multiple species were present in at least one study but did not repeat across studies; and (c) a cross-variance design when at least one species repeated across studies. In most cases the more complex structures (b-c) were required only due to a handful of species (i.e., the appropriate model for the majority of the dataset was a, but b or c were selected due to the presence of a handful of datapoints). In some of these cases, excessive model complexity led to non-converging and hence less reliable models. In such cases, the algorithm used a fallback model equivalent to the simple model (a). Model assumptions (normality and heteroscedasticity of the residuals; influential cases) were tested using qq-plots, scatterplots of the fitted vs. residual values, and leave-one-out. The full annotated script and all datafiles used in the analysis will be available upon publication at https://github.com/omernevo/data_analysis_FFT.

## RESULTS

### Flowers

For flowers, 8 out of 15 traits were either positively or negatively correlated with elevation, indicating a potential effect of temperature on the trait value (Figure 1, Table S7). Sepal and petal width, nectar volume, and floral longevity were all positively correlated with elevation. In contrast, sepal length, petal length, inflorescence length, and number of flowers were negatively correlated with elevation. (Figure 1, Table S7)

### Fruits

For fruits, patterns were similar to flowers despite a substantially smaller dataset and fewer functional traits for fruits represented. Four out of six functional traits showed a significant correlation, negative or positive, with elevation (Figure 2; Table S8). Vitamin C and crop size were positively correlated with elevation, indicating a potential decrease in both in warming environments. Weight and width showed significant negative correlations with elevation, indicating a potential increase in a warming world.

## DISCUSSION

Flowers and fruits form the interface of plant-animal interactions, driving plant reproduction and supporting many animal species. The presence, absence, and strength of interactions are governed by trait matching between plants and animals, and flower and fruit traits can be affected by environmental factors - among them temperature increase and other indirect effects of global climate change. Our objective was to assess whether temperature increase, proxied by an elevational gradient, systematically changes flower and fruit traits. We found consistent and significant effects in both flowers and fruits, indicating a potential effect of climate change on plant reproduction, animal communities, and ecosystem functioning.

### Flowers

In flowers, 8 out of 15 quantitative traits showed a significant systemic effect of elevation, forming three clusters: morphology, nutrient content, phenology & abundance. Measures of width (sepal, petal) were consistently positively correlated with elevation (i.e. predicted to decrease with warming temperatures), while measures of length (inflorescence, petal, sepal) were systematically negatively correlated with elevation (i.e. predicted to increase with rising temperatures). This indicates that within the normal range of those species, an increase in temperature is associated with flowers that are morphologically different, albeit not necessarily smaller or larger in size. These changes can reflect two mechanisms: a direct effect of temperature or other related exogenic factors leading to a change in the trait (Totland, 2001; Zhao and Wang, 2015; Plendl et al., 2024); or alternatively, a change of the flower to, for example, accommodate different pollinators across a gradient (Arroyo et al., 1982; Inouye and Pyke, 1988; Sun et al., 2014; Dellinger et al., 2021; Rabeau et al., 2026). Direct effects are more likely to disrupt trait-matching with pollinators, whereas indirect effects might reflect more adaptive (i.e. fitness-enhancing) responses that allow plants to adjust to changing pollinators. For example, increased bumble bee visitation in higher elevation has been shown to be associated with larger corollas due to increased attractiveness and fit (Maad et al., 2013). In this case, an indirect effect would be adaptive: environmental conditions such as temperature affect pollinator species and abundance, and flowers change in order to adapt to available pollinators. A mixture of both direct and indirect effects is possible: for example, the different directions of correlations between length and width traits could indicate a trade-off of maintaining visual attractiveness due to larger width while reduced length due to abiotic pressures.

Our results further indicate effects of temperature/elevation on flower longevity, longevity, and nectar volume. For reduced longevity in warming climates, this is generally congruent with patterns reported in multiple other studies (Blionis and Vokou, 2001; Fabbro and Korner, 2004; Pacheco et al., 2016; Arroyo et al., 2017) and argued to be a partial offset to slow pollination and improve fruit set (Arroyo et al., 2013, 2017). However, reduced longevity is not found in all alpine species (Arroyo et al., 2021). For the number of flowers and nectar volume the results reported are mixed and dependent on species and environment (Cruden, 1976; Mu et al., 2015). For example, in an alpine plant, experimental warming reduced nectar production by area up to 90% (Mu et al., 2015), in line with our results. From a plant perspective, these mixed patterns can be attributed to the same trade-offs described for size traits. From a pollinator perspective, changing temperature can result in temporal shifts in flower abundance (Aldridge et al., 2011) and such temperature induced changes in floral resources have had a strong impact on bee abundance (Ogilvie et al., 2017). Additionally, changes in flowering phenology due to early snowmelt or drought can further alter floral resources availability in montane regions as well (Pyke et al., 2016; Theobald et al., 2017; Kudo, 2020; Powers et al., 2022; Kudo et al., 2025). These changes might be especially detrimental for pollinator species, such as bumble bees, in higher elevation and colder climates given their adaptation to current systems.

However, the strong interconnectedness between plant and pollinators and context-specific responses of floral traits renders it difficult to predict the general impact of these changes on pollinators.

Taken together, these results raise the question how species may balance between maintenance of functional traits adapted to current pollinators and the need to potentially shift to ones whose abundance increases with temperature. Given the trait matching and co-variation between plants and pollinators (Egawa et al., 2020; Toji et al., 2021, 2022), the extent to which abiotic factors drive plants and pollinators into mismatching directions and limit the adaptation potential to the interaction patterns could be seen as key factors. Small changes in size can result in differences in pollination success (Nagano et al., 2014) and flower temperature can also affect pollinator visitations (Leal and Koski, 2023; Koski et al., 2024; Apland and Koski, 2025). Moreover, functional traits are often developmentally or mechanistically connected, meaning that they are unlikely to change independent of one another. For example, changes in morphology will lead to a change in the volume to surface area ratio, and hence potentially to changes in temperature; this can cascade to changes in microbial communities, volatilization of scent compounds and hence floral scent, water content, etc.

### Fruits

Elevation was significantly associated with multiple fruit functional traits, including shape (fruit width), size (mass), crop size, and nutrient content. A potential increase in fruit width will potentially make fruits less elongated and more round. This can primarily affect smaller birds that swallow fruits whole (“gulpers”), for whom a match between gape width and the dimensions of the fruit is critical (Wheelwright, 1985; Levey, 1987; Todeschini et al., 2020). This may be exacerbated by the fact that fruits as whole may become larger: fruit weight was also negatively correlated with elevation in our study - a result that is in line with previous studies (Fischer et al., 2022). Changes in fruit size and shape could also indirectly affect the nutritional value of the fruit: fruits generally hold large amounts of water and biomass, and their morphology is related to their ability to withhold the internal pressure from within (Valenta et al., 2022). A change in shape and mass can thus lead to a change in water content or connective tissue that is not nutritious but is required to maintain the integrity of the fruit. At the same time, it is important to note that fruit length, while not reaching statistical significance, showed a tendency in the same direction. As such, it remains possible that the absence of an effect on fruit length is a product of limited statistical power, in which case the effect on width would not reflect a change in fruit shape, but rather a general increase in fruit size. Nonetheless, the potential effects on both fruit-eating animals and physical integrity are still potentially applicable.

Crop size was positively correlated with elevation, indicating that it may decrease in a warming world. A decrease in crop size potentially indicates a less dense dispersal kernel of plants (Snell et al., 2019), and may negatively affect birds, who overall show preference for large crop sizes (Palacio and Ordano, 2018). This may not necessarily have negative implications to plant reproduction if the effect is consistent across species since competition for resources and germination sites may also be decreased. Yet it is unlikely that this effect is consistent across species, and it is therefore likely that this pattern may lead to an alteration of the plant community structures as it favors some species over others. Nonetheless, a decrease in fruit crop sizes is likely to be more consequential to animal communities: both frugivores (fruit eaters) and granivores (seed eaters) communities may decrease if fruit yields are systemically lower.

Finally, vitamin C (ascorbic acid) was also positively correlated with elevation, indicating a projected decrease in fruit vitamin C content in a warming world. Vitamin C is a critical micronutrient for vertebrates, but multiple lineages have lost the capacity to synthesize it internally. Many of those are fruit eaters and important seed dispersers: anthropoid primates (monkeys and apes), many bats, and passerine birds (Drouin et al., 2011). This is hardly surprising, as fruits are a key source of vitamin C, and its availability is likely to have relaxed selection pressures to retain autonomic synthesis in these lineages. As such, a decrease in vitamin C content can potentially negatively affect animal consumers. Indirectly, it may also affect plants as animals that are not obligate frugivores are forced to increase the fruit content in their diet. The dynamics here are complex, as this may on one hand lead to an increase in seed dispersal quantity, but a decrease in quality (Nevo et al., 2023).

### Synthesis

Flowers and fruits showed a surprisingly consistent response to elevated temperature across an elevational gradient in two domains: morphology (measures of width and length), and a decrease in nutritional value (nectar volume; vitamin C). In contrast, the number of flowers and fruits appear to go in opposite directions: flower number is projected to increase with temperature according to our results, but fruit crop is expected to drop. A decrease in longevity was found only for flowers, although it was not measured for fruits. This, however, may have a direct effect on fruit availability as the fruiting season may start earlier and hence also end earlier in the year. Indirectly, it may also be a driver of the decrease in fruit crop sizes, especially if sampling concentrated towards the end of the fruiting seasons of the included studies.

In general, the effects on fruits may be direct or indirect through flowers: fruits develop from flowers and pollination quality directly affects fruit functional traits (Wietzke et al., 2018).

Changes in flower traits are known to influence pollination success, potentially lowering the frequency and efficiency of visitors and thus decreasing fruit set. This may be the driver behind the higher proportion of fruit traits affected by climate change (66% of fruit traits, as opposed to 53% of flower traits), although the small number of traits makes this estimate prone to error. Our dataset was unfortunately too limited to disentangle the direct and indirect effects of temperature on fruit traits, but this can be tested using a smaller, more targeted, model system comparing the fruits of species in which flower traits do change to those that do not.

Alternatively, an analysis of fruit traits across temperature gradients of species that are abiotically-pollinated may provide an answer to the question whether the effects on fruits result directly from temperature effects, or whether they are the product of indirect effects due to changes in floral traits.

Given that flowers and fruits are developmentally linked, and that traits within each organ do not develop independent of one another, these results demonstrate the potential cascading effects of any change to such a complex system. Effective reproduction is a function of the development of fully-developed seeds, wrapped in a fruit whose functional traits are optimized for its most suitable local dispersers; which is on its own a function of high-quality pollination (Wietzke et al., 2018). Floral traits can be affected by temperature, or indirectly by the effect of temperature on other floral traits; and that the relevant fruit traits can be affected by temperature directly, other fruit traits, or pollination quality and hence floral traits. Thus, any change that may occur along the way may lead to measurable cascading effects on plant reproduction.

Regardless of the mechanism, the fact that both flowers and fruits are affected by temperature change is alarming. Given the prevalence of animal pollination (Tong et al., 2023) and seed dispersal, primarily in the tropics (Jordano, 2000), this means that many plant species will be affected, with potential negative consequences for the pollination success and seed dispersal. At the same time, it is critical to note that our analysis did not directly measure trait matching with animals. As such, it does not directly indicate that trait shifts within the ranges described here will necessarily substantially change the structure or strength of animal-plant interaction networks. Yet again, our work covered species in their natural habitats and in which variation is probably still within the “normal” range for the species. But if these patterns are consistent and will become more extreme as the planet warms, it is highly likely that some interactions will be lost, emerge, or change in intensity, at least for some communities. Further, it is possible that some of the changes we document are not necessarily a direct physiological effect of temperature on traits, but rather an adaptive response to different pollinator or seed disperser abundances. It is predicted that a major effect of climate change will be a shift in mutualist abundance (Arroyo et al., 2021; Guerra et al., 2025; Rabeau et al., 2026), which may even favor a shift in functional traits. At the same time, these shifts are unlikely to be tradeoff-free: different mutualists may provide pollination or dispersal services that vary in quantity and quality (Schupp, 1993; Nevo et al., 2023), and even a shift that is adaptive in the sense that it is optimized for a new normal may be suboptimal compared to previous conditions. As such, it is also highly likely that global warming will, if it has not already, drive shifts in animal-plant interaction networks affecting the majority of plant species and countless numbers of animal species and communities.

### Outlook

Our work demonstrates potential major disruptions to animal-plant interactions in a warming world. At the same time, it is important to highlight its limitations. First, while the elevation-for- temperature approach is commonly used to assess the effects of temperature on naturally occurring species, it cannot control for various confounding factors (Lovell et al., 2023).

Moreover, it mainly measures temperature, which is only the first immediate effect of climate change. A warming climate will experience several other shifts, in precipitation patterns, nutrient cycle, herbivory pressure, and many other factors (Harvey et al., 2023). Yet this unfortunate reality indicates that the effects we found represent a floor in the projected changes, not a ceiling; and that other factors will compound and drive further disruptions in animal-plant interactions.

Another factor is that while comprehensive, our analysis only covered traits that are widely measured across species and communities. These naturally include morphological traits that are easier to record, but lack in chemical traits. Floral and fruit scent (Raguso, 2008; Nevo et al., 2018a), non-volatile chemistry (Leonhardt et al., 2024; Nguyen et al., 2025), and nutrient content (Nevo et al., 2022; Barnett et al., 2023; Leonhardt et al., 2024) are major drivers of animal-plant interactions. But their analysis is complex, not fully comparable (Nguyen et al., 2026), and therefore all but absent from datasets in the numbers required for a meta analysis.

Yet another factor is that not all species and biomes are expected to respond in the same ways (Novaes et al., 2024), and that generally multiple confounding and interacting factors may affect the degree and even direction temperature affects fruits and flower traits. Our dataset was too limited to meaningfully parse it into different biomes or to control for other potential confounding factors. As such, our approach was to standardize the measures (z-transform) across all traits and elevations, under the assumption that the studied populations were sampled within their natural habitats and that by that our analyses simulates variation in temperatures realistic for each species and population. Nonetheless, future studies should aim at looking at the specific effects on each population under different circumstances.

### Conclusions

Flower and fruit traits are the interface of many animal-plant interactions and variation in them affects plant reproduction, animal communities, and most terrestrial ecosystems. Our work shows that increased temperature, approximated using the elevation-for-temperature approach, is associated with myriad changes in flower and fruit traits, affecting their morphology, availability, timing, and nutrient content. These imply that climate change is likely to have driven changes that may disrupt pollination and seed seed dispersal processes, and that these are likely to be exacerbated by future inevitable warming.

## ACKNOWLEDGMENTS

The project was funded by the iDiv Flexpool fund and the German Science Foundation (DFG; NE 2156/3-1). In memory of Geno (Eugene) Schupp, 1952-2026.

