## Supplementary material for "Effects of temperature gradient on flower and fruit traits: a meta analysis": Sup figs 1-2

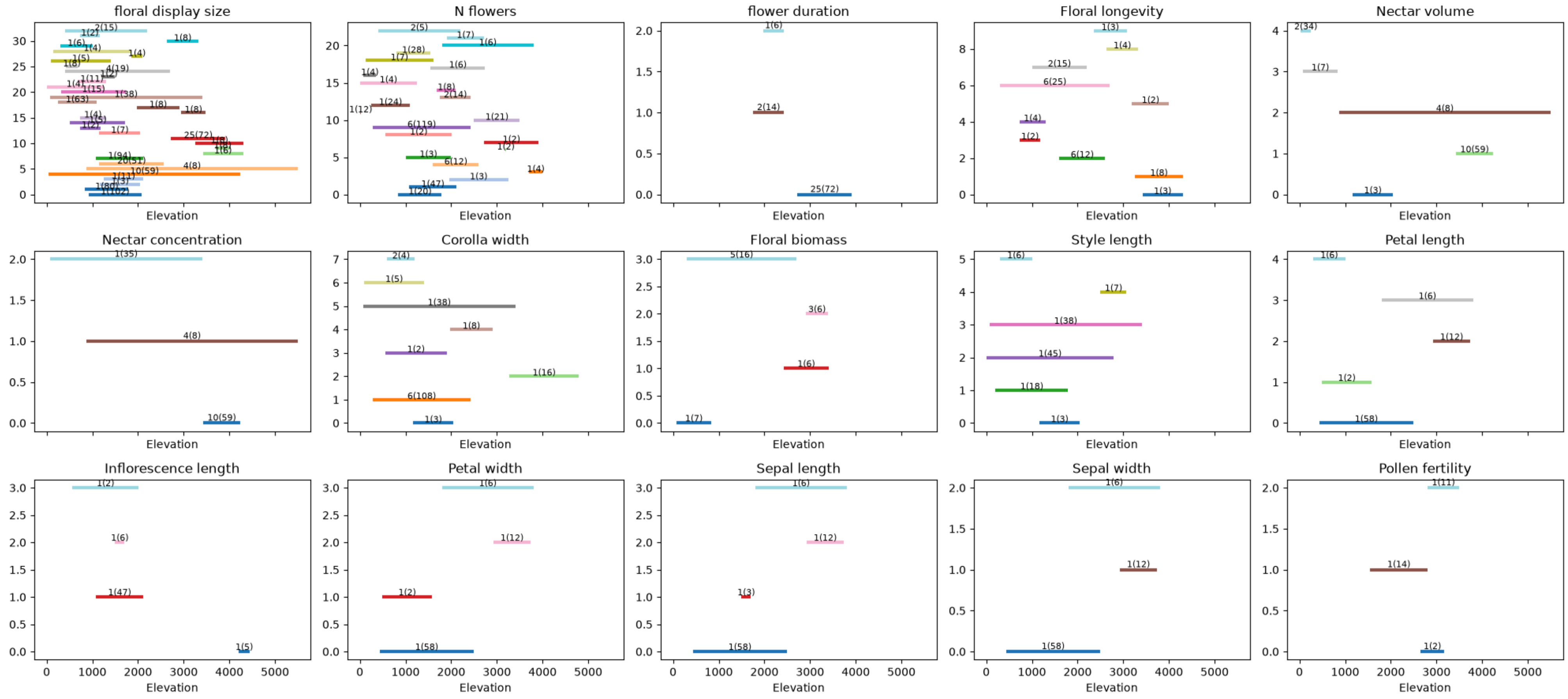


**Fig S1: Distribution of datapoints and studies across elevations for all floral traits included in the final analysis.** Each line is a study. Numbers represent the number of species within the study and total number of datapoints for all species in the study.

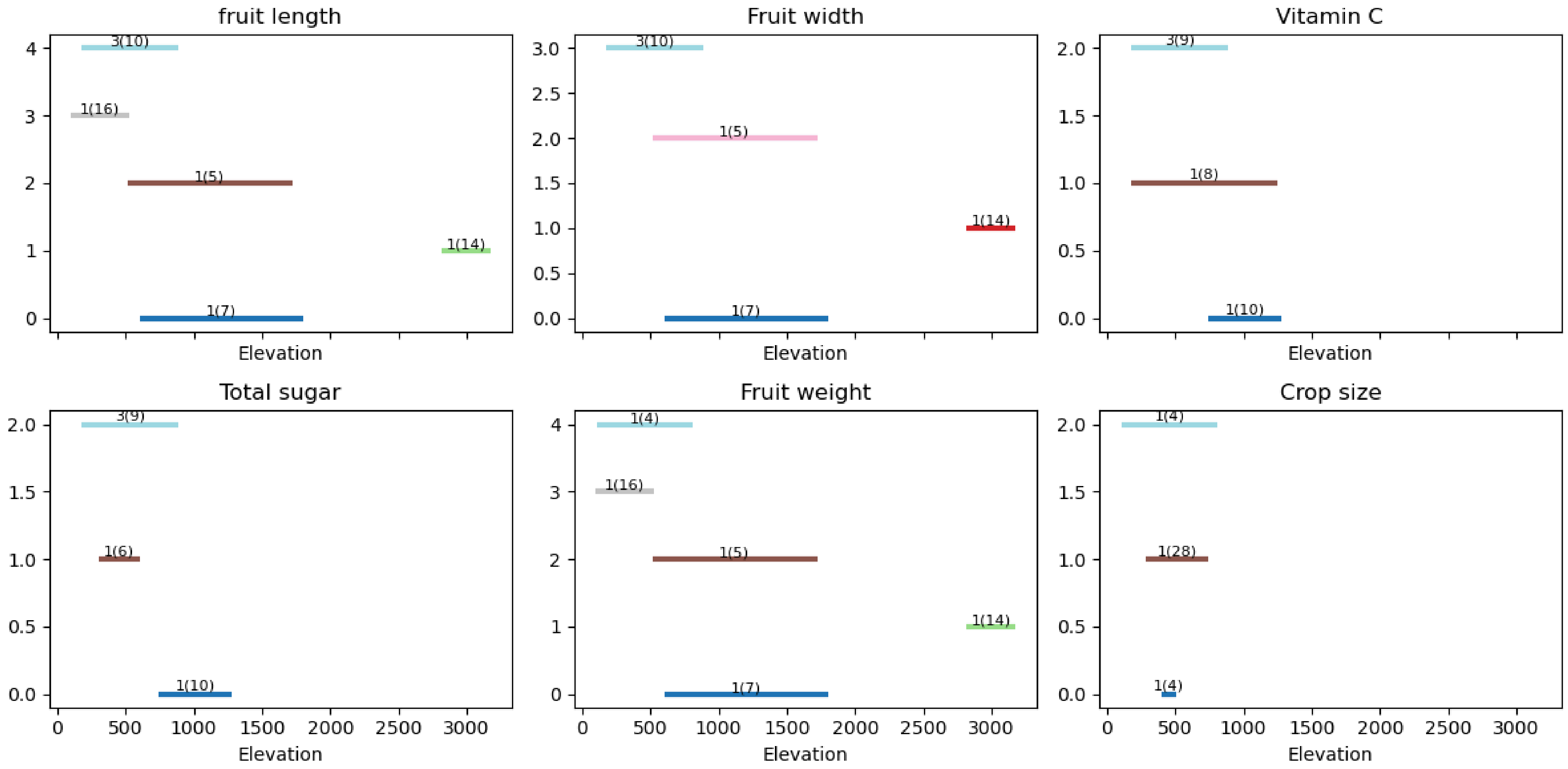


**Fig S2: Distribution of datapoints and studies across elevations for all fruit traits included in the final analysis.** Each line is a study. Numbers represent the number of species within the study and total number of datapoints for all species in the study.
